# Developmental prefrontal myelination shapes inhibitory interneuron function and adult social behavior

**DOI:** 10.64898/2026.10.06.757195

**Authors:** Takahira Yamauchi, Kohei Kamikawa, Kazuhiko Yamamuro, Hisayoshi Kubota, Minobu Ikehara, Tsutomu Takeda, Kazuki Okumura, Michihiro Toritsuka, Shouta Sugio, Atsuko Nagaoka, Mizuki Hino, Julio Leon, Satoshi Yoshinaga, Hiroki Ikeda, Kazuki Kurimoto, Yasuhiko Saito, Yasuto Kunii, Ken-ichiro Kubo, Hiroaki Wake, Taku Nagai, Manabu Makinodan

## Abstract

Early-life adversity and abnormal myelination are implicated in psychiatric disorders, but how developmental myelination influences adult social cognition remains unclear. Here, we tested whether disrupting developmental myelination in the medial prefrontal cortex (mPFC) impairs adult social behavior. Using complementary genetic strategies in male mice, we suppressed Myelin Regulatory Factor (*Myrf*)-dependent myelination locally in the juvenile mPFC or broadly in oligodendrocyte-lineage cells. Both approaches selectively impaired adult social novelty preference while sparing sociability, locomotion, and affective behaviors. Focal prefrontal dysmyelination also reduced social dominance. Spatial transcriptomics revealed coupling between myelin-related genes and parvalbumin (PV) interneuron markers in the healthy mPFC, which was disrupted by developmental hypomyelination. Consistent with this transcriptional alteration, PV interneurons showed reduced high-frequency firing despite preserved intrinsic excitability. In the human prefrontal cortex, MBP–PV expression was correlated in controls but not in individuals with schizophrenia. These findings indicate that developmental prefrontal myelination contributes to PV interneuron maturation and suggest that disrupted oligodendrocyte–interneuron coordination may contribute to impaired social recognition.

## Introduction

Adverse childhood experiences (neglect, abuse, and chronic stress) are risk factors for aberrant neurodevelopment and psychiatric disorders (Teicher *et al*., 2022; Teicher & Samson, 2016). Adversity during sensitive developmental periods can produce lasting changes in brain structure and function (Teicher *et al*., 2022; Teicher & Samson, 2016), particularly in regions with prolonged postnatal maturation such as the prefrontal cortex (PFC) (Selemon, 2013). Because the PFC remains immature and plastic through adolescence while supporting executive function, emotion regulation, and social cognition (Selemon, 2013), it may be especially vulnerable to environmental insults. Childhood maltreatment has been associated with reduced PFC volume, disrupted connectivity, emotion dysregulation, and impaired social functioning (McLaughlin *et al*., 2019).

Disrupted myelination may contribute to this developmental vulnerability. Myelination, the ensheathment of axons by oligodendrocyte-derived lipid membranes, enables rapid impulse conduction and temporal coordination of neural networks (Fields, 2008). Because PFC myelination continues through childhood and adolescence, early-life adversity may interfere with this process and contribute to persistent cognitive and emotional impairments (Teicher & Samson, 2016). Animal models show that early social stress alters PFC myelin development. Makinodan et al. identified a critical postnatal period during which social experience is required for normal oligodendrocyte maturation and prefrontal myelination; disrupted social interaction during this window produces persistent hypomyelination and later behavioral abnormalities (Makinodan *et al*., 2012). Similarly, Liu et al. showed that juvenile social isolation reduces PFC myelination, including myelin basic protein (MBP) expression and myelin thickness, and that these deficits persist after re-socialization (Liu *et al*., 2012).

Altered myelination is associated with psychiatric illness, including abnormal white matter integrity in the PFC and its connecting tracts in major depression, anxiety disorders, and schizophrenia (SCZ) (Cetin-Karayumak *et al*., 2020; Guo *et al*., 2024; Jenkins *et al*., 2016; Kelly *et al*., 2018; Shen *et al*., 2017; Tromp *et al*., 2012). Myelin also participates dynamically in experience-dependent plasticity: adaptive myelin remodeling supports motor learning, and activity-dependent myelin plasticity contributes to cortical synchronization (Kato *et al*., 2020; Xiao *et al*., 2016). Whether myelination causally regulates social behavior, however, remains unclear. Juvenile social isolation also compromises maturation of fast-spiking parvalbumin (PV)-positive interneurons, which regulate network synchronization and prefrontal-dependent social behaviors, including sociability and social hierarchy (Bicks *et al*., 2020; Zhang *et al*., 2021). Although transient juvenile demyelination impairs PV interneuron maturation in the medial prefrontal cortex (mPFC) of mice (Hijazi *et al*., 2025), how mPFC myelination interacts with PV interneurons to shape social behavior remains unresolved.

We used multiple genetic strategies to suppress Myelin Regulatory Factor (*Myrf*) during development and generate three mouse models of prefrontal hypomyelination. We tested whether impaired developmental mPFC myelination reproduces social abnormalities associated with social isolation and directly alters PV interneuron function.

## Materials and Methods

### Animals

All animal procedures were approved by the Animal Care Committee of Nara Medical University and conformed to the NIH Guide for the Care and Use of Laboratory Animals. Mice were housed under controlled temperature and humidity on a 12-hour light/dark cycle (lights on 08:00–20:00) with food and water ad libitum. Three complementary models were used to disrupt developmental myelination in the mPFC. *Myrf*^fl/fl^ mice (B6;129-*Myrf*^tm1Barr^/J; The Jackson Laboratory, Bar Harbor, ME, USA; stock #010607), carrying loxP sites flanking exon 8 of the *Myrf* gene, were crossed with tamoxifen-inducible *Olig2*-CreERT2 knock-in mice (B6.Cg-*Olig2*^tm1(cre/Esr1*)Htak^/Rbrc; RIKEN RBRC#01507). *Myrf*^fl/fl^;*Olig2*^wt/creER^ offspring were designated inducible conditional knockouts (cKO), with *Myrf*^fl/fl^;*Olig2*^wt/wtER^ littermates as controls. From P21 to P35, cKO and control littermates received tamoxifen-containing chow (0.5 mg/g normal chow TAM, Sigma Aldrich, Japan) *ad libitum* to induce recombination in Olig2-expressing oligodendrocyte precursor cells.

Reduced adult mPFC MBP immunoreactivity confirmed *Myrf* disruption. *Pvalb*-T2A-FlpO-D mice (B6.Cg-*Pvalb*^tm4.1(flpo)Hze^/J, JAX #022730) were crossed with *Myrf*^tm1Barr^ mice for Flp-dependent labeling of PV-positive interneurons.

Unless otherwise noted, behavioral and electrophysiological experiments used male mice to limit variability associated with estrous cycling. Transgenic strains were maintained on a C57BL/6J genetic background, and F1 hybrid males were used for mixed-background crosses to minimize genetic variability.

### Stereotaxic viral injections

For mPFC-specific *Myrf* deletion, P21 *Myrf*^fl/fl^ mice were anesthetized with isoflurane and placed in a stereotaxic frame (Narishige, Tokyo, Japan). AAV-DJ-CMV-eGFP-iCre (titer ∼1 × 10^13^ genome copies (GC)/mL; viral serotype DJ, cat# 7093, Vector Biolabs, Malvern, PA, USA) was delivered bilaterally to the prelimbic mPFC at four coordinates: (1) anteroposterior (AP)=+2.1 mm, mediolateral (ML)=+0.4 mm, dorsoventral (DV)=−2.5 mm; (2) AP=+1.9 mm, ML=+0.4 mm, DV=−2.3 mm; (3) AP=+2.1 mm, ML=−0.4 mm, DV=−2.3 mm; and (4) AP=+1.9 mm, ML=−0.4 mm, DV=−2.3 mm. In each hemisphere, 450 nL was injected at 60 nL/min, and the needle remained in place for 5 min to prevent backflow. Controls received an equivalent injection of AAV-DJ-CMV-eGFP (cat# 7101, Vector Biolabs). For electrophysiological labeling, *Myrf*^fl/fl^ mice were crossed with *Pvalb*-FlpO mice (B6.Cg-*Pvalb*^tm4.1(flpo)Hze^/J, JAX #022730), and a Flp-dependent reporter AAV (AAV-Ef1α-Coff/Fon-mCherry; Addgene, Watertown, MA, USA; plasmid #137134) was co-injected with AAV-Cre (or control AAV) into the mPFC of these mice. In PV^FlpO^ mice, this reporter virus induces robust expression of mCherry in PV interneurons, enabling their visualization, except in cells co-expressing Cre (in which the “Cre-off” cassette suppresses mCherry). This strategy produced regional oligodendrocyte *Myrf* deletion while labeling PV interneurons in the same mPFC region for targeted recording. All AAV vectors were commercially prepared and validated for genomic titer and purity. Mice were monitored daily after surgery and recovered for at least 4 weeks before behavioral testing or tissue analysis.

For focal demyelination of the mPFC via MBP translation knockdown, we used an AAV serotype 8 vector carrying the 3’ untranslated region of *MBP* mRNA (AAV8-MBP3’UTR), which impairs *MBP* mRNA transport/translation. P21 C57BL/6J male mice received bilateral mPFC injections of AAV8-MBP3’UTR (∼1 × 10^12^ GC/mL, 0.5 µL per side at 0.1 µL/min); controls received equal-titer AAV8-CMV-GFP. Stereotaxic coordinates and injection parameters were identical to those used for AAV-Cre. Mice matured to 8–10 weeks before behavioral testing. Reduced MBP staining in the injected mPFC confirmed focal demyelination, whereas control mice showed normal myelin patterns.

### Behavioral tests

Behavioral testing was performed in adult mice (2–3 months old) during the light phase. Mice were habituated to the testing room for at least 30 min, and experimenters were blinded to genotype/group during testing and analysis.

### Three-chamber social interaction test

Sociability and social novelty preference were assessed in a previously described three-chamber apparatus (Komori *et al*., 2024), a plexiglass box (60 × 40 cm^2^) with three interconnected chambers (Ohara & Co., Ltd., Tokyo, Japan). Mice were habituated to the empty apparatus for 10 min the day before testing.

On the test day, mice were placed in the center chamber and allowed to explore all three chambers containing two empty wire mesh cylinders for 10 min.

The cylinders were positioned in the side chambers to acclimate the mouse to their presence. For the sociability phase (Session 1), an unfamiliar age- and sex-matched C57BL/6J mouse was enclosed in one cylinder, and an identical empty cylinder (object) was placed in the opposite chamber. The subject explored for 10 min while chamber and sniffing times were recorded using ANY-maze (Stoelting Co., Wood Dale, IL, USA). Sociability was quantified as the social preference index (time with mouse − time with object).

After a brief interval, the social novelty phase (Session 2) began. The familiar mouse from Session 1 remained in its cylinder, and a novel mouse was placed in the opposite chamber. After 10 min of exploration, social novelty preference was calculated as time with the novel mouse − time with the familiar mouse.

### Augmented Reality-Based Long-Term Animal Behavior Observation (AR-LABO) social assay

Social behavior was also assessed in a group setting using AR-LABO, an automated platform for continuous tracking of freely interacting mice in a home-cage–like environment. At least 1 week before testing, each mouse received an identification (ID) tag printed with ArUco markers, attached dorsally under anesthesia with an elastic string, and treated with quinine to prevent removal or biting by cagemates. One subject and three unfamiliar age- and sex-matched C57BL/6J mice were placed in a single cage (276 × 445 × 204 mm^3^; CL-0128; CLEA Japan Inc., Tokyo, Japan) and monitored by infrared camera at 20 frames per second. Videos were analyzed offline using our custom tracking system, now commercially available from O’Hara & Co., Ltd.

The center of each ID tag was recorded as X–Y coordinates, exported as comma-separated values (.csv) files, and processed in R (version 4.2.1) (Endo *et al*., 2021; Endo *et al*., 2019; Endo *et al*., 2018; Komori *et al*., 2024).

Coordinates exceeding 1.1 times the estimated cage floor area were excluded. Misidentified ID tag positions with Z-scores >3, calculated within a 2-s window (40 frames) before and after each time point, were removed. Missing coordinates in individual frames were linearly interpolated from adjacent valid detections.

We used a two-session paradigm to examine sociability and social novelty. In Session 1 (sociability), the subject interacted freely for 60 min in a clean cage with three unfamiliar male C57BL/6J partner mice. After Session 1, the subject was briefly removed and two of the three partners were replaced with unfamiliar mice, leaving one familiar partner. The subject was then returned for a 10-min Session 2 (social novelty) with the familiar and novel mice. AR-LABO recorded approach behavior throughout both sessions.

Social approach was defined as one mouse coming within ∼3 cm of another. Position data were used to calculate approaches made and received by the subject toward novel and familiar mice. Cages were cleaned and new stimulus mice were used for each subject to minimize olfactory carryover.

Locomotor activity was quantified as the total distance traveled by each mouse’s ID tag. A contact event was defined when two ID-tag coordinates were ≤20 mm apart. For each contact, distances traveled by both mice during the preceding 1 s (20 frames) were compared; the mouse traveling farther was classified as approaching and the other as receiving. Approach and receive counts were summed for each mouse.

### Open-field test

For the open-field test, each mouse was placed in the center of a white plastic chamber (40 × 40 × 40 cm^3^) and allowed to explore for 10 min. ANY-maze tracked total distance traveled and time spent in the central zone as measures of locomotor activity and anxiety-like behavior, respectively.

### Tail-suspension test (TST)

The TST was performed as previously described (Yin *et al*., 2011). Mice were suspended by adhesive tape placed approximately 2 cm from the tail tip, with the head ∼15 cm above the apparatus base. Behavior was recorded for 8 min, and the final 5 min were analyzed using ANY-maze. Immobility, defined as absence of movement except respiration, was quantified during this 5-min period.

### IntelliCage

The IntelliCage system (TSE Systems GmbH, Bad Homburg, Germany) is a computerized, fully automated apparatus for studying spontaneous emotional and cognitive behaviors in group-housed radio-frequency identification (RFID)-tagged mice (Kubota *et al*., 2025; Lipp *et al*., 2024). The system consists of a home cage (55 × 37.5 × 20.5 cm^3^) containing four triangular operant corners (15 × 15 × 21 cm^3^) and can concurrently monitor up to 16 RFID-tagged mice. Each corner is accessible to only one mouse because of the dimensions of the tunnel and corner and is equipped with RFID readers, infrared sensors, two nose-poke holes, lickometers, and motorized gates controlling access to water bottle nipples. The system automatically records corner visits, nose pokes, licks, mouse ID, and corner ID. Behavioral testing comprised an adaptation phase followed by a social dominance test. During the first 3 days of adaptation, doors remained open to permit free water access. During the subsequent 6 days, mice opened a door by nose poke to obtain 5-s water access, with each door opening only once per corner visit. Visits, nose pokes, and licks were monitored continuously, and circadian activity was analyzed during the final 3 days. For the social dominance test, mice were habituated to water restriction for 7 days. They were deprived of water for 21 h/day and allowed 5-s access to water per corner visit from 20:15 to 23:15. A red LED and speaker provided visual and auditory cues during the 3-h session. On day 7, corner visit duration during the first 10 min and across the full 3-h session was recorded to assess competition for limited water rewards.

### Immunohistochemistry

Free-floating coronal sections containing the mPFC were processed for MBP immunofluorescence. Sections were permeabilized by washing in phosphate-buffered saline (PBS) with 0.3% Triton X-100 (PBS-Tx), incubated in 25 mM glycine/PBS to quench residual aldehydes, treated with 0.3% H₂O₂ in methanol, and blocked for 30 min at room temperature in PBS-Tx containing 5% normal goat serum (NGS). Sections were then incubated overnight at 4°C with two mouse monoclonal anti-MBP antibodies (clone SMI-94, IgG1; clone SMI-99, IgG2b; BioLegend, San Diego, CA, USA), diluted in PBS-Tx with 1% NGS and applied simultaneously. After washing, sections were incubated for 1 h at room temperature with Alexa Fluor 546-conjugated goat anti-mouse IgG (1:1000), washed, rinsed in Tris buffer (50 mM Tris– HCl, pH 7.4), mounted, and imaged by epifluorescence microscopy (BZ-X700, Keyence, Osaka, Japan). For MBP3’UTR-injected mice, immunohistochemistry followed a previously described protocol (Kato *et al*., 2020). Mice were anesthetized and transcardially perfused with ice-cold PBS followed by 4% paraformaldehyde. Brains were post-fixed overnight at 4°C, cryoprotected overnight in PBS containing 30% sucrose, and sectioned coronally at 30 μm using a sliding microtome (Leica SM2000R, Wetzlar, Germany). Sections were washed in PBS containing 0.3% Triton X-100, blocked for 1 h at room temperature in 5% NGS (Vector Laboratories, Newark, CA, USA, #S-1000-20), and incubated overnight at 4°C with rat anti-MBP antibody (Millipore, Billerica, MA, USA, #MAB386). After washing, Alexa Fluor 488 goat anti-rat IgG (Invitrogen, Waltham, MA, USA; #A11006, diluted 1:1,000) was applied for 1 h at room temperature. Sections were then mounted and imaged by fluorescence microscopy (BZ-9000, Keyence). MBP immunoreactivity in the mPFC was quantified using ImageJ 1.52k (NIH, Bethesda, MD, USA), and the mean of three sections per mouse was used for statistical analysis.

### Electrophysiology

Brain slices containing the mPFC were prepared from 61-day-old mice generated by crossing *Pvalb*-T2A-FlpO-D (*Pvalb*-2A-FlpO-D) mice with *Myrf*^tm1Barr^ mice. At postnatal day 21 (P21), mice received bilateral mPFC injections of pAAV-Ef1a-Coff/Fon-mCherry together with either AAV-DJ-eGFP-iCre (Cre⁺) or AAV-DJ-CMV-eGFP (Cre⁻), enabling Flp-dependent PV interneuron labeling and Cre-dependent regional *Myrf* deletion. Mice were deeply anesthetized with isoflurane and decapitated, and brains were rapidly transferred to ice-cold (approximately 4°C) sucrose-based cutting solution bubbled with 95% O₂/5% CO₂ and containing (in mM): 210.3 sucrose, 2.5 KCl, 26.2 NaHCO₃, 1.0 NaH₂PO₄, 0.5 CaCl₂, 4.0 MgSO₄, and 11.0 D-glucose. Coronal mPFC slices (300 µm) were prepared using a vibrating tissue slicer (Vibratome 1000 Plus 102; Pelco International, Redding, CA, USA) and incubated for at least 60 min in standard artificial cerebrospinal fluid (ACSF) containing (in mM): 119.0 NaCl, 2.5 KCl, 26.2 NaHCO₃, 1.0 NaH₂PO₄, 2.0 CaCl₂, 2.0 MgCl₂, and 11.0 D-glucose, maintained at 32°C and bubbled with 95% O₂/5% CO₂. Slices were then held in ACSF at room temperature (approximately 25°C). During recording, slices were superfused with ACSF at 32°C and 2 mL/min. mCherry-positive PV interneurons were identified by epifluorescence and targeted under infrared differential interference contrast optics (BX51WI; Olympus, Tokyo, Japan). Fluorescence images were obtained using an X-Cite XYLIS LED light source (model XT720S; Excelitas Technologies, Pittsburgh, PA, USA), a U-FMCHE fluorescence filter cube (Evident Scientific, Tokyo, Japan), and a DP23M digital microscope camera (Evident Scientific). Whole-cell current-clamp recordings were performed using a Multiclamp 700A amplifier (Axon Instruments, Molecular Devices, San Jose, CA, USA). Patch pipettes (3–5 MΩ) contained (in mM): 127.5 K-methanesulfonate, 5.0 KCl, 2.0 MgCl₂, 2.0 Mg-ATP, 0.3 Na-GTP, 0.6 ethylene glycol tetraacetic acid (EGTA), and 10.0 HEPES, with pH adjusted to 7.25 using KOH. Membrane potentials were corrected for the liquid junction potential (10 mV for Cs-based solutions). Data acquisition and stimulation were controlled using Signal 4 software with a Power 1401 interface (Cambridge Electronic Design, Cambridge, UK).

### Current-clamp recording

During current-clamp recordings, series resistance was monitored and compensated using a bridge circuit, and pipette capacitance was corrected. Voltage signals were low-pass filtered at 10 kHz and digitized at 20 kHz. The baseline membrane potential was maintained near −70 mV via current injection. Action potential (AP) analysis was performed using Signal 4 software (version 4, Cambridge Electronic Design). The AP threshold was defined as the membrane potential at which the first derivative of the voltage trace (dV/dt) exceeded 10 mV/ms during a depolarizing current step. The AP amplitude was measured as the voltage difference between the threshold and the peak of the first spike evoked by a 100-pA current injection. To assess APs and subthreshold membrane properties, we recorded membrane potential responses to hyperpolarizing and depolarizing current pulses (500 ms duration). Depolarizing pulses ranging from 10–200 pA were applied in 10 pA increments.

### Voltage-clamp recording

During voltage-clamp recordings, pipette capacitance was compensated, and series resistance was continuously monitored but not compensated. Only recordings with a stable series resistance ≤20 MΩ were included in the analyses. Current signals were low-pass filtered at 800 Hz and digitized at a sampling frequency of 10 kHz. Spontaneous excitatory postsynaptic currents (EPSCs) and spontaneous inhibitory postsynaptic currents (IPSCs) were recorded in standard ACSF. EPSCs were recorded at a holding potential of −70 mV to isolate excitatory currents, and IPSCs were recorded at 0 mV to isolate inhibitory currents. Spontaneous EPSCs and IPSCs were analyzed using Mini Analysis software (Synaptosoft Inc., Decatur, GA, USA). Events were detected using an amplitude threshold of 5 pA. A 10-min stable segment of continuous recording was selected for analysis.

### Quantitative real-time PCR (qRT-PCR)

qRT–PCR was performed using the DRaqL (direct RNA recovery and quenching for laser capture microdissection) method (Ikeda *et al*., 2023; Ikeda *et al*., 2024; Shishido *et al*., 2023). For laser microdissection (LMD), mouse brains were rapidly removed, frozen on dry ice, stored at −80°C, sectioned coronally at 25 μm through the mPFC, and mounted on 2.0-μm polyethylene naphthalate (PEN) membrane slides (Leica Microsystems; Cat. No. 11505189). After brief dehydration, the mPFC was selectively isolated using a LEICA LMD7000 according to the manufacturer’s instructions. Before dissection, a 200 μL microcentrifuge tube cap was loaded with 6.4 µL ice-cold cell lysis buffer containing RNase inhibitor and ERCC RNA Spike-In Mix (final dilution 1:500,000). Dissected tissue was collected by gravity into the buffer on the tube cap. After each collection, the tube was immediately closed, tapped, and briefly centrifuged (5,000 × *g*, 10 s) to recover the lysate and minimize sample loss from evaporation. Samples remained on ice throughout the procedure. For cDNA synthesis and amplification, lysates were incubated at 72°C for 6 min to promote tissue lysis and RNA denaturation and then cooled on ice. Deoxycholate (DOC) was neutralized with 2.8 μL quenching buffer (2.5× SuperScript II buffer, 25% Triton X-100, and 5 mg/mL bovine serum albumin), followed by 1.6 μL template-switching RT mixture (6.25 μM N-template-switching oligo, 31.25 U each of SuperScript II and SuperScript III, and 5 U recombinant RNase inhibitor). RT was performed using the programmed cycle for full-length cDNA synthesis. Subsequently, 15 μL PCR master mix (1.67× SeqAmp buffer, 0.33 μM N-IS PCR primer, and 0.5 μL SeqAmp DNA polymerase) was added (final reaction volume 25.8 μL), and cDNA was amplified using a SeqAmp-specific program. Amplified cDNA was purified using NucleoMag NGS Clean-up and Size Select magnetic beads (Macherey-Nagel, Düren, Germany) with 0.8× bead volume, two 80% ethanol washes, and elution in 50 μL nuclease-free water, then stored at −20°C. qRT-PCR was performed on a QuantStudio 6 Flex Real-Time PCR System (Applied Biosystems, Waltham, MA, USA) using TB Green™ Premix Ex Taq™ II (Tli RNaseH Plus) (Takara Bio, Shiga, Japan; Cat. No. RR820A), gene-specific primers, and purified cDNA. Each 10 μL reaction contained 5 μL TB Green Premix, 0.4 μM each of forward and ERCC reverse primers, and 1 μL cDNA. Cycling conditions were 95°C for 30 s; 40 cycles of 95°C for 5 s and 60°C for 30 s; and melt-curve analysis at 95°C for 15 s, 60°C for 1 min, then heating to 95°C at 0.05°C/s. Reactions were performed in technical triplicate. Gene expression was normalized using RPL19 as an endogenous reference and ERCC RNA as an external spike-in control. Relative expression was calculated by the ΔΔCt method and expressed as fold change relative to control samples. Primer sequences are listed in Supplementary Table 1.

### Spatial transcriptomics of adult mouse brain

Visium analysis was performed using the adult AAV-mediated mPFC-restricted *Myrf* deletion model. Brains were rapidly removed, snap-frozen, and sectioned coronally (10 μm) to include the mPFC. Sections were mounted on Visium Spatial Gene Expression slides (10x Genomics, Pleasanton, CA, USA) and processed according to the manufacturer’s protocol. After hematoxylin and eosin staining and imaging, spatially barcoded libraries were prepared using the Visium Spatial Gene Expression 3’ kit and sequenced on an Illumina platform (Illumina, San Diego, CA, USA). Space Ranger generated spot-level gene expression matrices aligned to histological images. Analyses were restricted to GFP-positive spots within the viral target region, where the relationship between MBP and PV expression was evaluated using correlation analyses.

### Spatial transcriptomics of adult human brain

Postmortem brain samples from 5 individuals with SCZ and 4 control individuals were obtained from the Tohoku Postmortem Brain Bank for Psychiatric Research (https://www.fmu-bb.jp/english/index.htm) (Leon *et al*., 2024; Miyahara *et al*., 2023). This study was approved by the ethics committees of Fukushima Medical University, the Tohoku University Graduate School of Medicine, RIKEN Yokohama Campus, Keio University School of Medicine, and The Jikei University School of Medicine. All procedures were performed after obtaining written informed consent from the next of kin. SCZ diagnoses were based on criteria from the fourth or fifth edition of the Diagnostic and Statistical Manual of Mental Disorders (DSM-IV or DSM-5). The daily dose of antipsychotics for the 3 individuals with SCZ during the 3 months before death was expressed as the chlorpromazine-equivalent dose (CP-eq). The anterior cingulate cortex (approximately 10 × 10 × 10 mm^3^, Brodmann area 24) was identified in frozen coronal slabs according to a human brain atlas. To assess the correlation between *MBP* and *PVALB* expression in postmortem brains from individuals with SCZ and matched controls, we analyzed the Visium spatial transcriptomic dataset, which we previously reported (Mooijman *et al*., 2016). Spatial transcriptomic reads were aligned to the human reference genome (hg38) using Space Ranger (v2.0.1, 10x Genomics). Data processing was performed in R (v4.1.2) using Seurat (v4.1), applying spot-level filtering thresholds of 350–5,000 detected features, mitochondrial gene content <40%, and hemoglobin gene content <20%. We performed unbiased clustering from 30 Harmony-corrected principal components. Clusters were annotated based on their spatial context and expression of canonical marker genes.

### Statistical analysis

Statistical analyses were performed using GraphPad Prism 9.3.1. All tests were two-sided, with p<0.05 considered statistically significant. Parametric tests were used when assumptions of normality and homoscedasticity were met; otherwise, non-parametric tests were applied. Data are presented as mean ± standard error of the mean (SEM) unless otherwise stated, with exact p values and sample sizes in the figure legends. Independent two-group comparisons used two-tailed unpaired *t*-tests or, when parametric assumptions were violated, two-sided Mann–Whitney U tests. Within-group comparisons in the three-chamber social interaction test used two-sided paired *t*-tests. Spearman’s rank correlations assessed MBP–PV spatial associations in mouse and human datasets and the relationship between MBP intensity and competitive performance in the IntelliCage task. Electrophysiological input–output relationships were analyzed using two-way repeated-measures analysis of variance (ANOVA) with current as the within-subject factor and genotype as the between-subject factor. Human MBP–PV correlations were analyzed within clusters corresponding to cortical layers 2/3 and 5/6. The numbers of animals, cells, and spatial spots analyzed are indicated in the corresponding figure legends.

## Materials availability

All unique materials generated in this study are available from the corresponding author upon reasonable request.

## Results

### Developmental mPFC myelination is required for adult social novelty preference

Early-life social isolation reduces mPFC myelination and impairs adult social behavior (Liu *et al*., 2012; Makinodan *et al*., 2012), but whether these effects are causally related is unclear. To test whether developmental mPFC myelination is required for adult social behavior, we bilaterally injected *Myrf*^fl/fl^ (*Myrf*^tm1Barr^) mice at postnatal day 21 (P21) with AAV/DJ-GFP-iCre (Cre⁺) or control AAV/DJ-CMV-eGFP (Cre⁻) (Figure 1a). In adults, Cre⁺ mice showed reduced *Myrf* mRNA relative to Cre⁻ controls (Figure 1b) and decreased MBP immunoreactivity in the mPFC (Figure 1c), confirming region-specific hypomyelination. We first assessed behavior using the three-chamber social interaction test. During the sociability phase, both groups spent significantly more time exploring the social target than the object, indicating intact sociability (Figure 1d). During social novelty testing, however, Cre⁻ mice preferred the novel mouse whereas Cre⁺ mice did not (Figure 1e), indicating selective impairment of social novelty preference. Because the three-chamber test permits only indirect social contact and has limited ecological validity (Fukumitsu *et al*., 2022; Jabarin *et al*., 2022), we next used AR–LABO to assess freely interactive social behavior (Figure 1f). In Session 1 (60 min), social approaches and locomotor activity did not differ between groups (Figure 1g). In Session 2 (10 min), Cre⁺ mice made significantly fewer approaches toward the novel mouse than Cre⁻ controls, while locomotor activity remained comparable (Figure 1h). Open-field and TST outcomes also did not differ between groups, indicating no detectable change in anxiety- or depression-like behavior (Figure 1—figure supplement 1a, b). Together, these findings show that early postnatal *Myrf* deletion in the mPFC selectively impaired adult social novelty preference while preserving general sociability and locomotor activity.

**Figure 1.**
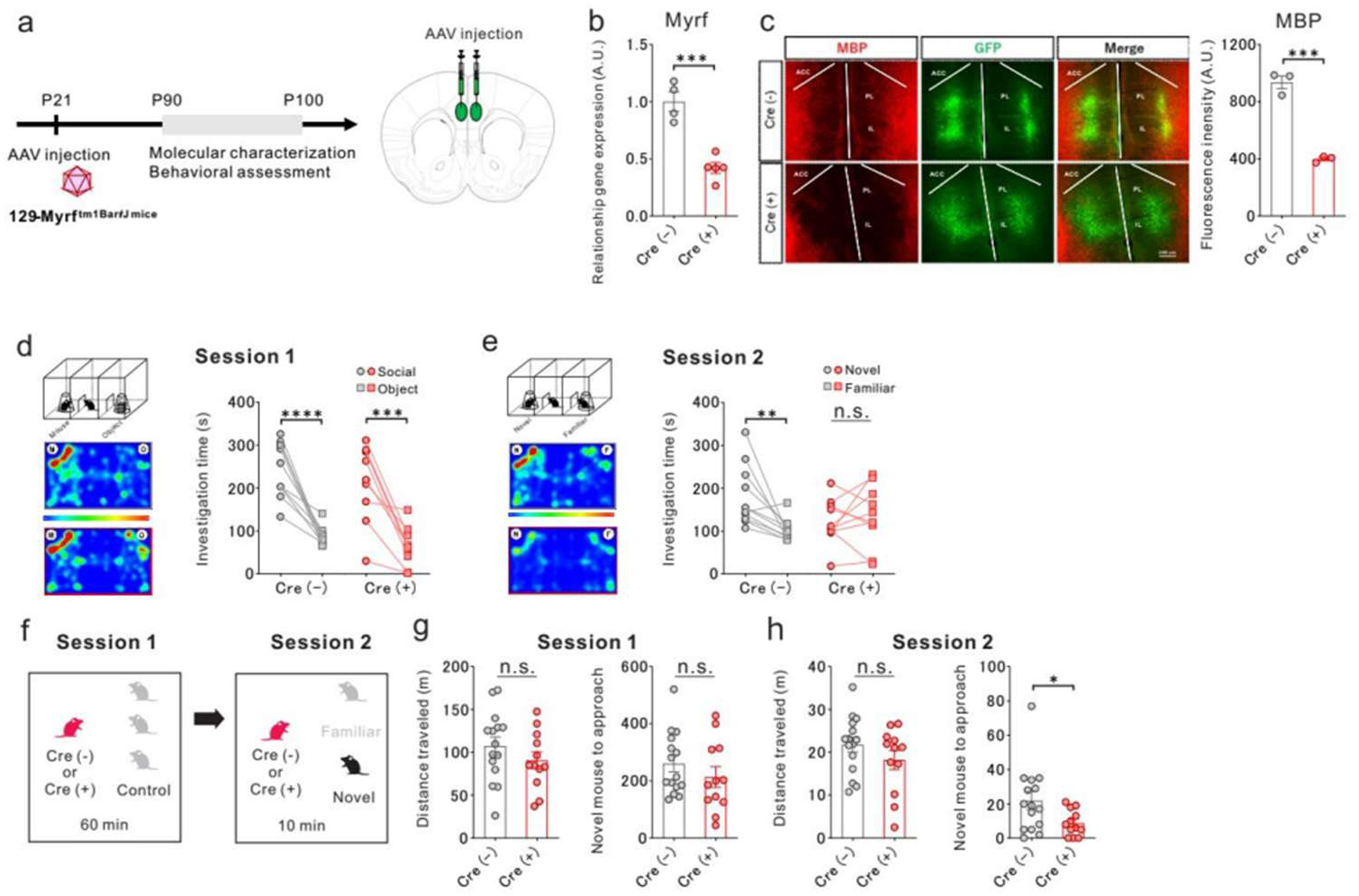
Developmental medial prefrontal cortex (mPFC) myelination is required for adult social novelty preference. (a) Experimental design showing bilateral injection of AAV/DJ-GFP-iCre (Cre⁺) or control AAV/DJ-CMV-eGFP (Cre⁻) into the mPFC of *Myrf*^fl/fl^ mice at postnatal day 21 (P21). (b) Quantitative RT–PCR analysis showing reduced *Myrf* mRNA expression in the mPFC of Cre⁺ mice compared with Cre⁻ controls. (c) Representative images and quantification of myelin basic protein (MBP) immunoreactivity showing a significant reduction in myelin density in the mPFC of Cre⁺ mice compared with Cre⁻ controls (two-tailed unpaired t-test, p=0.0004; Cre⁻, n=3; Cre⁺, n=3), confirming region-specific hypomyelination. Scale bar=200 µm. (d, e) Three-chamber social interaction test. (d) Sociability phase (Session 1). Quantification of investigation time spent with a mouse (Social) versus an object (Object). Both Cre⁻ and Cre⁺ mice spent significantly more time investigating the mouse than the object (two-sided paired t-test; Cre⁻, p<0.0001, n=11 pairs; Cre⁺, p=0.0002, n=10 pairs). (e) Social novelty phase (Session 2). Quantification of investigation time spent with a novel mouse (Novel) versus a familiar mouse (Familiar). Cre⁻ mice showed a significant preference for the novel mouse, whereas Cre⁺ mice did not (two-sided paired t-test; Cre⁻, p=0.0059, n=11 pairs; Cre⁺, p=0.5333, n=10 pairs), indicating a selective impairment in social novelty preference in Cre⁺ mice. (f) Schematic of the Augmented Reality-Based Long-Term Animal Behavior Observation (AR–LABO) system used to assess freely interacting social behavior. (g, h) AR–LABO assay. (g) Session 1 (sociability phase). Total locomotor activity did not differ between Cre⁻ and Cre⁺ mice (left panel; two-sided unpaired t-test, p=0.2627; Cre⁻, n=15; Cre⁺, n=12). The number of social approaches toward other mice was also comparable between groups (right panel; two-sided unpaired t-test, p=0.3031; Cre⁻, n=15; Cre⁺, n=12). (h) Session 2 (social novelty phase). Total locomotor activity during Session 2 did not differ between the groups (left panel; two-sided unpaired t-test, p=0.2089; Cre⁻, n=15; Cre⁺, n=12). In contrast, the number of approaches toward novel mice was significantly reduced in Cre⁺ mice compared with Cre⁻ controls (right panel; two-sided Mann–Whitney U test, p=0.0475; Cre⁻, n=15; Cre⁺, n=12). AAV, adeno-associated virus; A.U., arbitrary units; GFP, green fluorescent protein; eGFP, enhanced green fluorescent protein; RT-PCR, reverse transcription polymerase chain reaction; n.s., not significant.

### Developmental *Myrf* deletion in Olig2-lineage cells recapitulates social novelty deficits

To independently validate the role of developmental myelination in social behavior, we crossed B6;129-*Myrf*^tm1Barr^/J mice with B6.Cg-*Olig2*^tm1(cre/Esr1*)Htak^/Rbrc mice to generate inducible conditional knockout (cKO) mice. *Myrf*^fl/fl^;*Olig2*^wt/creER^ cKO mice and *Myrf*^fl/fl^;*Olig2*^wt/wt^ littermate controls (CTR) received tamoxifen-containing chow from P21 to P35, inducing *Myrf* deletion in Olig2-expressing cells during developmental myelination (Figure 2a). Adult cKO mice showed reduced mPFC MBP immunoreactivity and significantly lower MBP signal intensity than controls, confirming disruption of developmental myelination (Figure 2b).

**Figure 2.**
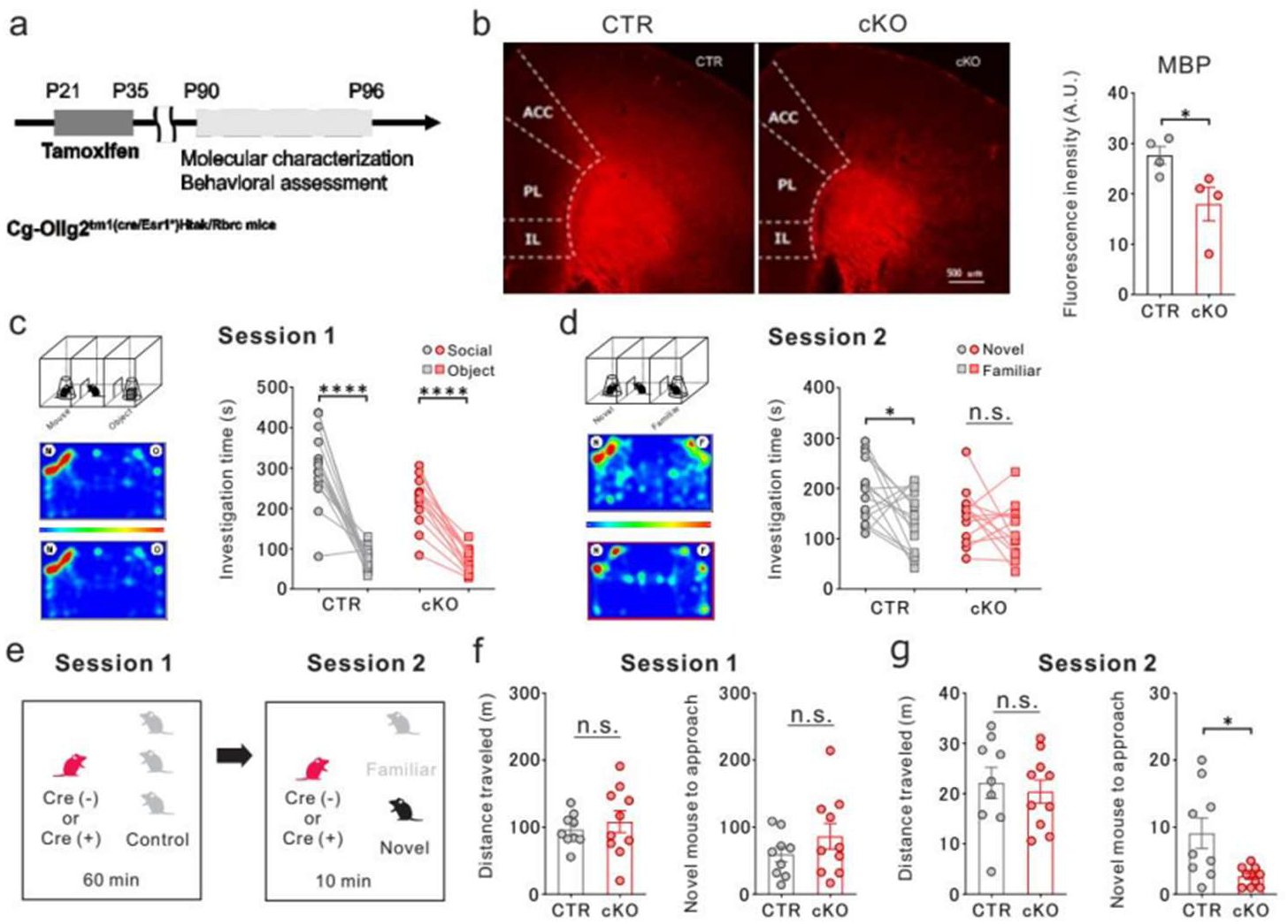
Developmental *Myrf* deletion in Olig2-lineage cells recapitulates social novelty deficits. (a) Experimental timeline illustrating tamoxifen administration (P21–P35) in *Myrf*^fl/fl^;*Olig2*^wt/creER^ (conditional knockout, cKO) and littermate control (*Myrf*^fl/fl^;*Olig2*^wt/wt^, CTR) mice. (b) Representative images and quantification of myelin basic protein (MBP) immunoreactivity showing a significant reduction in myelin density in the medial prefrontal cortex (mPFC) of cKO mice compared with CTR mice (two-sided unpaired t-test, p=0.0431; CTR, n=4; cKO, n=4). Scale bar=500 µm. (c, d) Three-chamber social interaction test. (c) Sociability phase (Session 1). Quantification of investigation time spent with a mouse (Social) versus an object (Object). Both CTR and cKO mice spent significantly more time investigating the mouse than the object (two-sided paired t-test; CTR, p<0.0001, n=15 pairs; cKO, p<0.0001, n=13 pairs). (d) Social novelty phase (Session 2). Quantification of investigation time spent with a novel mouse (Novel) versus a familiar mouse (Familiar). CTR mice showed a significant preference for the novel mouse, whereas cKO mice did not (two-sided paired t-test; CTR, p=0.0159, n=15 pairs; cKO, p=0.2041, n=13 pairs), indicating a selective impairment in social novelty preference in cKO mice. (e) Schematic of the Augmented Reality–Based Long-Term Animal Behavior Observation (AR– LABO) system used to assess freely interacting social behavior. (f, g) AR–LABO assay. (f) Session 1 (sociability phase). Total locomotor activity did not differ between CTR and cKO mice (left panel; two-sided unpaired t-test, p=0.2542; CTR, n=9; cKO, n=10). The number of social approaches toward other mice was also comparable between groups (right panel; two-sided unpaired t-test, p=0.5415; CTR, n=9; cKO, n=10). (g) Session 2 (social novelty phase). Total locomotor activity during Session 2 did not differ between the groups (left panel; two-sided unpaired t-test, p=0.6575; CTR, n=9; cKO, n=10). In contrast, the number of approaches toward novel mice was significantly reduced in cKO mice compared with CTR controls (right panel; two-sided unpaired t-test, p=0.0101; CTR, n=9; cKO, n=10). A.U., arbitrary units; CTR, control; n.s., not significant.

The cKO model recapitulated the mPFC-specific AAV–Cre phenotype. In the three-chamber sociability phase, both cKO and control mice preferred the mouse to the object (Figure 2c), whereas only controls preferred the novel mouse during the social novelty phase (Figure 2d). AR–LABO testing yielded similar results (Figure 2e): social approaches and locomotor activity did not differ during Session 1 (Figure 2f), but cKO mice made fewer approaches toward the novel mouse during Session 2 while locomotor activity remained unchanged (Figure 2g). Open-field and TST outcomes also did not differ between groups (Figure 2—figure supplement 1a, b). Thus, developmental *Myrf* deletion in Olig2-lineage cells selectively impaired social novelty preference without altering general sociability, activity, or anxiety- and depression-like behavior.

### Developmental prefrontal hypomyelination impairs social dominance behavior

To examine the role of developmental prefrontal myelination in social dominance, we used the IntelliCage system, which continuously assesses activity and social behavior in group-housed mice (Fig. S4a). To establish focal prefrontal hypomyelination, AAV8 carrying the MBP translation-inhibitory sequence (MBP3’UTR) under the myelin-associated glycoprotein (MAG) promoter was injected into the mPFC at P21, and behavioral testing began at P63 (Figure 3a). MBP3’UTR-injected mice showed reduced MBP immunoreactivity across cortical layers I–V and significantly lower mPFC MBP signal intensity than control AAV-injected littermates (Figure 3b, c; Figure 3—figure supplement 1a, b), confirming effective MBP suppression in the targeted region. During adaptation, circadian visit, nose-poke, and lick patterns over 24-h cycles were comparable between groups (Figure 3—figure supplement 2b–d), indicating preserved baseline locomotor activity, motivation, and task learning. After adaptation, mice underwent 7 days of water restriction, with water available as a reward during a 3-h session (20:15–23:15) to assess social competition (Figure 3a, d). MBP3’UTR-injected mice had markedly shorter visit durations in the competition corner during the first 10 min (Figure 3e), but this difference was not evident across the full 3-h session (Figure 3f). Total visit counts and overall locomotor activity remained comparable between groups. MBP signal intensity across cortical layers I–V correlated positively with corner visit duration during the first 10 min (Figure 3g). These findings associate focal mPFC hypomyelination with reduced competitive behavior under limited water access without evidence of generalized motor or motivational impairment.

**Figure 3.**
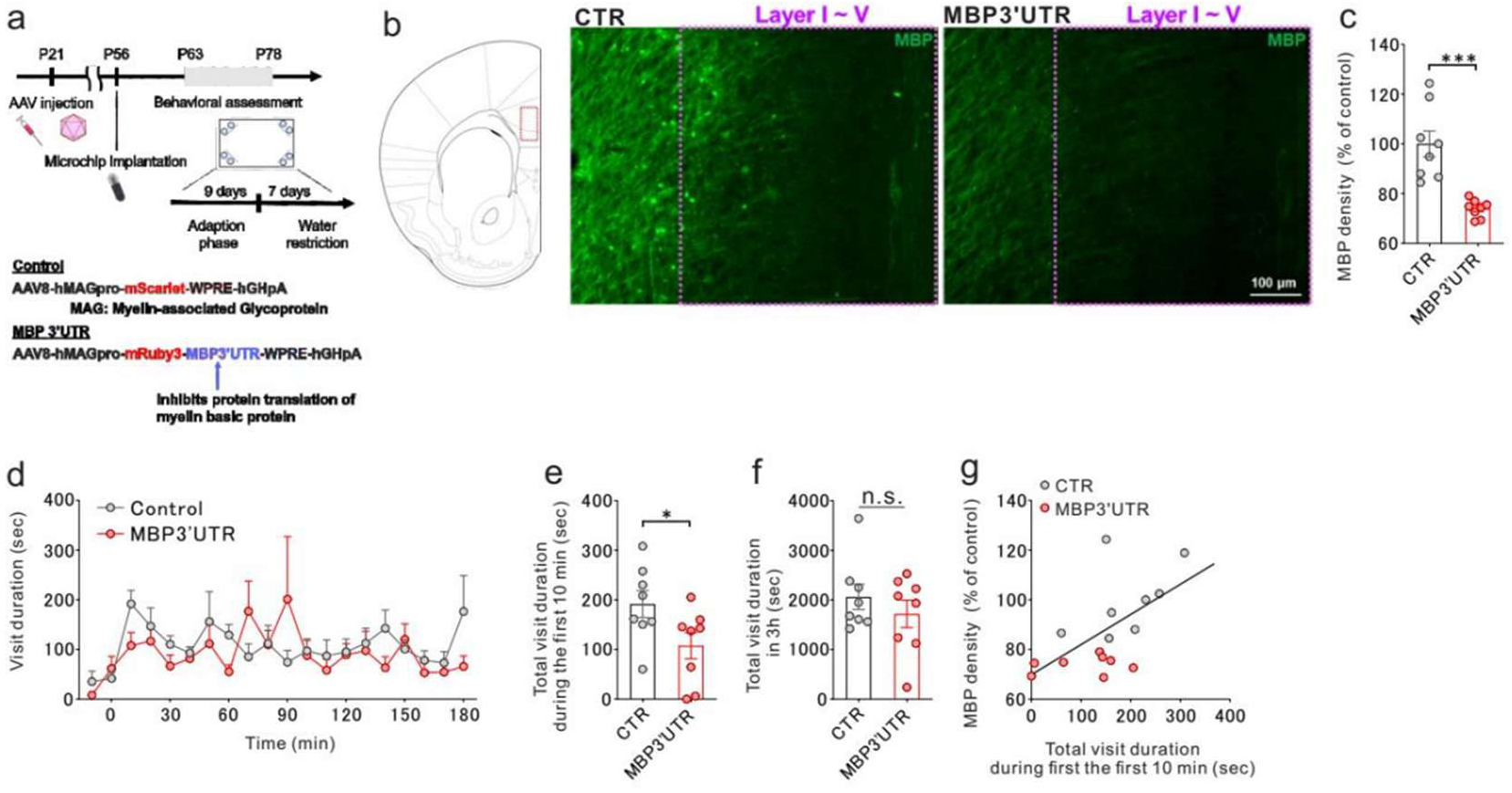
Prefrontal hypomyelination impairs social dominance behavior in the IntelliCage system. (a) Schematic of AAV8-mediated suppression of myelin basic protein (MBP) translation in the medial prefrontal cortex (mPFC). An AAV8 vector carrying the MBP translation-inhibitory sequence (MBP3’UTR) under the control of the myelin-associated glycoprotein (MAG) promoter was bilaterally injected into the mPFC at postnatal day 21 (P21). The experimental design for IntelliCage-based social dominance testing is shown. Following a 9-day adaptation period and 7 days of water restriction, mice competed for limited water access during a 3-hour daily session (20:15–23:15). The IntelliCage system enables continuous automated monitoring of individual behavior under group-housed conditions using radio-frequency identification (RFID)-based corner access control (see Figure 3—figure supplement 2a). (b) Representative immunofluorescence images of MBP immunoreactivity in the mPFC. The MBP signal was markedly reduced across cortical layers I–V in MBP3’UTR-injected mice (right) compared with control littermates (left). Scale bar=100 µm. (c) Quantification of MBP fluorescence intensity in the mPFC. MBP signal intensity was significantly decreased in hypomyelinated mice relative to controls, confirming effective regional reduction in MBP signal (two-sided unpaired t-test, p<=0.0001; Control, n=15; MBP3’UTR, n=14). (d) Time course of visit duration in the IntelliCage social competition assay. Visit duration was plotted across the 3-h testing period for control and MBP3’UTR-expressing mice. (e) Total visit duration during the first 10 min of the social competition session. MBP3’UTR-injected mice exhibited significantly shorter visit durations during the initial 10 min than control AAV-injected mice (two-sided unpaired t-test, p=0.0249; Control, n=15; MBP3’UTR, n=14). (f) Total corner visit duration across the full 3-hour competition period. No significant difference was observed between groups (two-sided unpaired t-test, p=0.5472; Control, n=15; MBP3’UTR, n=14), with no group difference over the full competition period. (g) Correlation between prefrontal MBP intensity and competitive performance. MBP signal intensity across cortical layers I–V showed a significant positive correlation with mean corner visit duration during the initial 10 min of the competition task (Pearson correlation, r=0.5135, p=0.0044, n=29), indicating that prefrontal MBP signal intensity is associated with early-phase competitive behavior under resource-limited conditions. AAV, adeno-associated virus; MBP3’UTR, myelin basic protein 3′ untranslated region; CTR, control; n.s., not significant.

### Developmental mPFC hypomyelination disrupts PV interneuron transcriptional and functional integrity

To identify cell types affected by developmental hypomyelination, we performed Visium spatial transcriptomics of the mPFC in *Myrf*^fl/fl^ (*Myrf*^tm1Barr^) mice injected with AAV/DJ-GFP-iCre (Cre⁺) or control AAV/DJ-CMV-eGFP (Cre⁻) (Figure 4a). In Cre⁻ mice, MBP, a canonical myelin marker, showed a robust positive spatial correlation with PV, a marker of fast-spiking GABAergic interneurons (Figure 4b). Regions with higher MBP expression also showed higher PV expression, whereas this MBP–PV relationship was markedly attenuated in Cre⁺ mice (Figure 4b), indicating disruption of their normal spatial coupling. qRT–PCR confirmed reduced *Pvalb* mRNA in the mPFC after AAV-mediated *Myrf* deletion (Figure 4—figure supplement 1a). In contrast, GAD67 and GAD65 mRNA levels were unchanged (Figure 4—figure supplement 1b, c), suggesting a preferential PV-related rather than generalized inhibitory-neuron effect.

**Figure 4.**
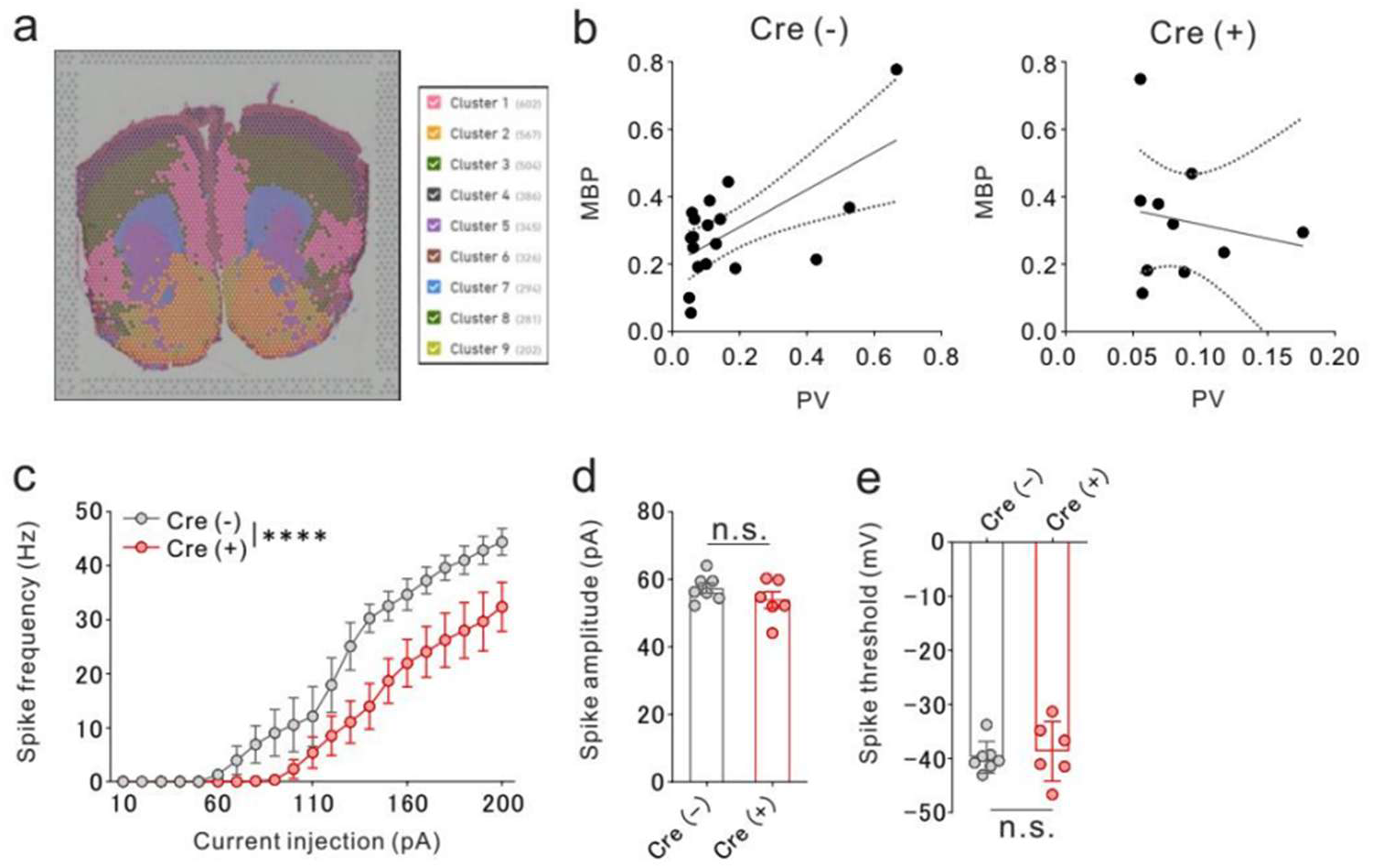
Developmental hypomyelination in the medial prefrontal cortex (mPFC) disrupts myelin–PV spatial coupling and PV interneuron firing. (a) Spatial transcriptomic (Visium) analysis of the mPFC showing spatial expression maps of the myelin marker myelin basic protein (MBP) and parvalbumin interneuron marker PV. (b) Scatter plot depicting a significant positive spatial correlation between MBP and PV expression in Cre⁻ (control) mice (Spearman’s ρ=0.510, p=0.031, n=18), which was not detected in Cre⁺ mice (Spearman’s ρ=−0.213, p=0.552, n=10). (c) Whole-cell current-clamp recordings from mCherry-labeled PV interneurons in the mPFC following region-specific *Myrf* deletion. Input–output curves of PV interneurons in Cre⁻ and Cre⁺ mice. Two-way repeated-measures analysis of variance (ANOVA) revealed a significant current × genotype interaction (F(19,209) =3.08, p=3.8 × 10⁻⁴), a significant main effect of current (F(2.78,30.62) =79.40, p<3.1×10⁻¹⁴), and a significant main effect of genotype (F(1,11) =6.53, p=0.027). n=7 cells from 4 biologically independent Cre⁻ mice and n=6 cells from 4 biologically independent Cre⁺ mice. (d) Spike threshold of PV interneurons did not differ between Cre⁺ and Cre⁻ mice (two-tailed unpaired t-test, p=0.658; n = 7 cells from 4 biologically independent Cre⁻ mice, n = 6 cells from 4 biologically independent Cre⁺ mice). (e) Spike amplitude was comparable between the Cre⁺ and Cre⁻ PV interneurons (two-tailed unpaired t-test, p=0.226; n = 7 cells from 4 biologically independent Cre⁻ mice, n = 6 cells from 4 biologically independent Cre⁺ mice). Cre−, control group; Cre+, *Myrf*-deletion group; n.s., not significant.

Developmental hypomyelination also did not produce overt microglial activation: CD11b, CX3CR1, and CD68 expression remained comparable to controls (Figure 4—figure supplement 1d–f), providing no evidence of a broad microglial response. We next assessed whether altered MBP–PV coupling was accompanied by changes in PV interneuron physiology. B6;129-*Myrf*^tm1Barr^/J (JAX 010607) mice were crossed with B6.Cg-*Pvalb*^tm4.1(flpo)Hze^/J (JAX 022730) mice, and at P21 the mPFC was bilaterally injected with pAAV-Ef1a-Coff/Fon-mCherry (Addgene #137134) plus either AAV-DJ-eGFP-iCre (Addgene #7093; Cre⁺) or AAV-DJ-CMV-eGFP (Addgene #7101; Cre⁻), enabling regional *Myrf* deletion while labeling PV interneurons for targeted electrophysiology. Whole-cell current-clamp recordings from mCherry-labeled PV interneurons showed significant main effects of injected current (F(2.784,30.62) =79.40, p<3.1 × 10⁻¹⁴) and genotype (F(1, 11) =6.53, p=0.0267), together with a significant current × genotype interaction (F(19,209) =3.08, p=3.8 × 10⁻⁴) (Figure 4c). Thus, the current-dependent firing response differed between Cre^⁺^ and Cre^⁻^ mice, with lower firing output under local hypomyelination. In contrast, basic AP properties were preserved: spike amplitude (t(11) =1.28, p=0.226) and threshold (t(11) =0.455, p=0.658) did not differ between groups (Figure 4d, e), and variance was comparable for both parameters (F-test, p>0.15). These results indicate that region-specific *Myrf* deletion reduces the firing capacity of PV interneurons without detectable changes in the basic biophysical properties of AP generation.

### Spatial transcriptomics of human postmortem brains shows MBP–PV correlation in controls but not in SCZ

To assess the translational relevance of the MBP–PV interneuron association observed in our animal model, we examined postmortem human anterior cingulate cortex using spatial transcriptomics. Visium profiling was performed in individuals with SCZ and matched controls, and we evaluated the correlation between *MBP* and PV expression levels. In control individuals, *MBP* and PV mRNA expression were significantly positively correlated (Figure 5a), indicating that regions with higher myelin-marker expression tended to show higher PV interneuron-marker expression. In contrast, this correlation was not detected in individuals with SCZ (Figure 5b), suggesting disruption of the normal spatial relationship between myelin- and PV-related expression. These human data parallel our animal findings, with MBP–PV correlation detected in controls but not in SCZ.

**Figure 5.**
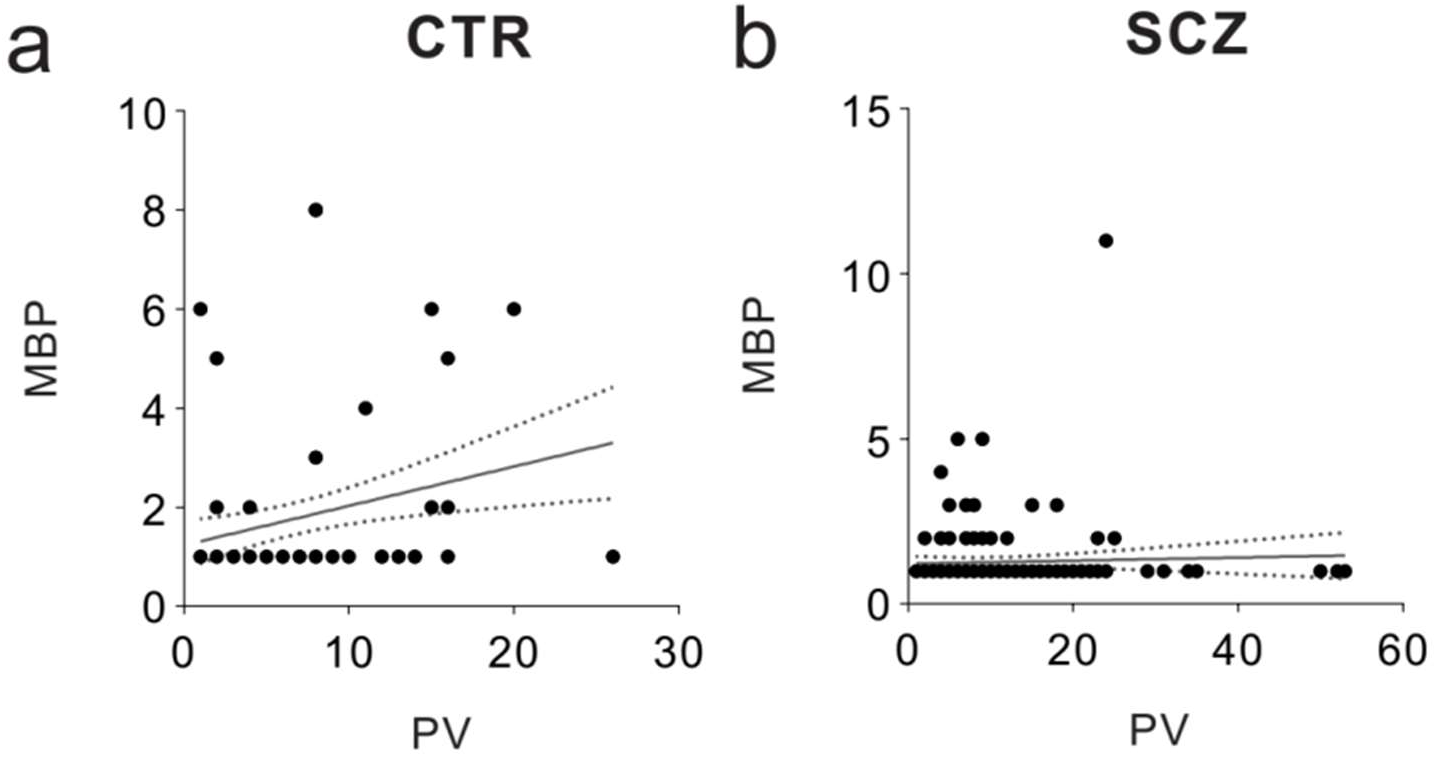
Spatial transcriptomic analysis of postmortem human prefrontal cortex shows a significant myelin basic protein (MBP)–parvalbumin (PV) correlation in controls but not in individuals with schizophrenia (SCZ). (a) Control group (postmortem prefrontal cortex [PFC] from n=4 individuals). Scatter plot showing a significant positive correlation between MBP and parvalbumin expression levels across spatial spots (Spearman’s r=0.27, p=0.0056; n=102 spatial spots after quality filtering). (b) SCZ group (postmortem PFC from n=5 individuals). Scatter plot showing no significant correlation between MBP and PV expression (Spearman’s r=0.01, p=0.916; n=171 spatial spots). CTR, control.

## Discussion

In this study, complementary genetic approaches showed that disrupting *Myrf*-dependent myelination during a postnatal developmental period, either locally in the mPFC or across oligodendrocyte-lineage cells, selectively impaired adult social novelty preference while sparing general sociability, locomotion, and affective behaviors. Developmental hypomyelination also disrupted spatial MBP–PV coupling and reduced high-frequency firing of PV interneurons despite preserved basic AP properties, consistent with previous work (Hijazi *et al*., 2025). Human postmortem spatial transcriptomics similarly showed disrupted myelin–PV coupling in the anterior cingulate cortex of individuals with SCZ. Together, these findings support a role for developmental prefrontal myelination in PV interneuron functional maturation and link oligodendrocyte–interneuron coordination to social cognitive function.

Developmental mPFC myelination may organize inhibitory circuit maturation beyond its established role in facilitating axonal conduction. This interpretation is supported by evidence that inhibitory-neuron axons, particularly those of PV interneurons, are myelinated, making myelin an important structural feature of fast-spiking inhibitory circuits rather than an exclusive property of long-range projection pathways (Micheva *et al*., 2016; Stedehouder *et al*., 2017). Consistent with this view, PV interneuron axon myelination directly regulates inhibitory circuit function (Benamer *et al*., 2020).

Early-life social deprivation produces lasting prefrontal hypomyelination and behavioral abnormalities, supporting a postnatal period in which myelin formation is experience-sensitive (Liu *et al*., 2012; Makinodan *et al*., 2012). Here, disrupting *Myrf*-dependent myelination during this period altered PV interneurons at molecular and physiological levels. In control mPFC, myelin-related gene expression was spatially coupled to PV interneuron markers (Hijazi *et al*., 2025), whereas developmental hypomyelination disrupted this relationship. These findings suggest that appropriate myelination supports oligodendrocyte–PV interneuron coordination and PV interneuron function during development.

Cortical myelination is increasingly recognized as an experience- and activity-sensitive process that can shape circuit development and behavior (McKenzie *et al*., 2014; Xiao *et al*., 2016).

Developmental studies have identified a postnatal critical period in which social experience regulates prefrontal myelination and long-term cognitive outcomes (Liu *et al*., 2012; Makinodan *et al*., 2012), supporting a developmentally instructive role for myelin. Adult learning studies similarly show that oligodendrocyte-lineage engagement, including *Myrf*-dependent oligodendrocyte differentiation and myelin programs, is required for skill acquisition (McKenzie *et al*., 2014; Xiao *et al*., 2016). Mechanistically, oligodendrocyte-lineage cells receive synaptic input from neurons and participate in bidirectional signaling (Bergles *et al*., 2000; Fang *et al*., 2022), providing a substrate through which circuit activity can influence myelin architecture and myelin can, in turn, stabilize circuit function. Cortical myelin is not distributed uniformly across axon classes; substantial myelination occurs on local inhibitory interneuron axons, particularly those of fast-spiking PV cells (Micheva *et al*., 2016; Mitew *et al*., 2018; Stedehouder *et al*., 2017). Large-scale anatomical and cell-type–resolved studies further indicate that individual oligodendrocytes can show a bias toward inhibitory targets. Activity-dependent myelin remodeling has also been demonstrated on PV interneuron axons, linking interneuron activity to changes in myelination patterns (Mitew *et al*., 2018; Stedehouder *et al*., 2018). Loss of PV-axon myelination compromises fast inhibition and circuit synchronization, including oscillatory dynamics (Dubey *et al*., 2022), supporting a functional role for PV myelination in cortical computation. Our findings therefore align with the emerging “myelin–PV unit” concept in neuropsychiatric disease models emphasizing interneuron timing and network coordination (Marín, 2016).

Our findings extend literature emphasizing conduction speed and synchrony as primary outputs of myelination (McKenzie *et al*., 2014; Xiao *et al*., 2016) by supporting a model in which developmental mPFC myelination contributes to PV circuit maturation across molecular, physiological, and behavioral levels. PV interneuron pathology in psychiatric disorders is often attributed to cell loss or transcriptional downregulation (Hashimoto *et al*., 2003), but postmortem evidence indicates that PV deficits in SCZ may reflect reduced molecular marker expression rather than neuronal loss (Lewis *et al*., 2003). Similarly, primary oligodendrocyte impairment can produce PV interneuron dysfunction without significant neuronal loss (Borges *et al*., 2023). Together, these observations are consistent with developmental myelination acting upstream of PV interneuron dysfunction, although the present data do not establish this pathway directly.

The selective deficit in social novelty preference with preserved baseline sociability is relevant to social cognitive dysfunction in psychiatric disorders. Social cognition is an important determinant of real-world functional outcome in SCZ and mediates the relationship between neurocognition and daily functioning (Fett *et al*., 2011; Green *et al*., 2015). Meta-analytic evidence indicates impairments across multiple domains of social cognition in SCZ (Savla *et al*., 2013), and medial prefrontal systems are implicated in social-cognitive processing (Amodio & Frith, 2006; Cao *et al*., 2018; Mack *et al*., 2024). In rodents, social novelty preference is commonly used to assess social recognition memory and social salience, processes modulated by prefrontal circuits (Chen *et al*., 2024; Sakamoto & Yashima, 2022). Preserved sociability with impaired novelty preference therefore suggests altered higher-order social information processing rather than a nonspecific reduction in social motivation. Consistent with this pattern, impaired communication between oligodendrocyte precursor cells and PV interneurons in the mPFC selectively disrupts social novelty preference (Fang *et al*., 2022). Interneuron hypomyelination has also been associated with higher-order behavioral deficits, including cognitive inflexibility, in rodent SCZ models (Maas *et al*., 2020). Our findings thus link developmental oligodendrocyte/myelin perturbation and PV interneuron dysfunction with later social recognition deficits.

A major strength of this study is that the disrupted myelin–PV relationship observed in mice was also detected in human postmortem tissue. Visium profiling showed positive spatial coupling between *Mbp* and *Pvalb* mRNA in the mPFC of control mice and between *MBP* and *PVALB* mRNA in the anterior cingulate cortex of control individuals. In mice, this spatial association is consistent with a relationship between local myelin-related expression and PV interneuron development, although spatial transcriptomics cannot establish whether individual PV axons are myelinated. The human finding is consistent with transcriptomic and imaging studies implicating both myelin and PV interneuron abnormalities in SCZ (Cetin-Karayumak *et al*., 2020; Hakak *et al*., 2001; Kelly *et al*., 2018; Lewis *et al*., 2005; Tasic *et al*., 2018; Tkachev *et al*., 2003; Uhlhaas & Singer, 2010) and with the proposed convergence of PV interneuron myelination in SCZ pathophysiology (Stedehouder & Kushner, 2016). Individuals with SCZ and myelin deficits have also been reported to show more severe social abnormalities than those with relatively preserved myelin (Pérez-Iglesias *et al*., 2010; Voineskos *et al*., 2013). Recent age-stratified transcriptomic analyses further suggest that earlier-onset SCZ, which is associated clinically with more severe symptoms and poorer outcomes, is accompanied by greater reductions in PV interneuron signatures (Kiss *et al*., 2026). Direct evidence linking early-onset SCZ to human myelination deficits remains limited; however, developmental studies suggest that disruption of adolescent myelination could affect PV interneuron maturation. In our study, spatial myelin–PV correlations were not detected in either hypomyelinated mice or individuals with SCZ. Because this loss of correlation may arise through multiple mechanisms, including compensatory changes in PV interneuron development, mechanistic interpretation requires further study.

This study has some limitations. First, although the multi-model approach supports a causal role for developmental myelination in behavioral outcomes, whether PV interneuron dysfunction mediates these effects remains unresolved. Second, we targeted a presumed postnatal critical period of prefrontal myelination without directly comparing disruption across developmental stages; therefore, the temporal specificity and reversibility of these effects remain to be determined. Third, the human postmortem sample was small, and these findings should be considered preliminary.

Together, our findings support a model in which developmental myelination of the mPFC contributes to inhibitory circuit maturation. Disruption of this program produces oligodendrocyte– PV uncoupling and selective impairment of social recognition. By linking developmental myelination with inhibitory circuit dysfunction across cell types, these results provide a mechanistic framework for understanding later social cognitive phenotypes.

## Acknowledgments

We thank the families of the deceased for their donations of brain tissue and for the time and effort devoted to the consent process and interviews. We also thank Ms. H. Onuma and Ms. M. Sawada for their contribution to coordinating the donations. We thank 10X Genomics and ACGT Core for providing reagents and processing some of the human samples.

## Author Contributions

Takahira Yamauchi: Conceptualization, Data curation, Formal analysis, Investigation, Methodology, Visualization, Writing – original draft, Writing – review and editing

Kohei Kamikawa: Conceptualization, Data curation, Formal analysis, Investigation, Methodology, Visualization

Kazuhiko Yamamuro: Investigation, Methodology, Formal analysis

Hisayoshi Kubota: Investigation, Methodology, Formal analysis

Minobu Ikehara: Investigation, Methodology, Formal analysis

Tsutomu Takeda: Investigation, Methodology, Formal analysis

Kazuki Okumura: Investigation, Methodology, Formal analysis

Michihiro Toritsuka: Investigation, Methodology, Formal analysis

Shouta Sugio: Investigation, Methodology, Formal analysis

Atsuko Nagaoka: Investigation, Methodology, Formal analysis

Mizuki Hino: Investigation, Methodology, Formal analysis

Julio Leon: Investigation, Methodology, Formal analysis

Satoshi Yoshinaga: Investigation, Methodology, Formal analysis

Hiroki Ikeda: Investigation, Methodology, Formal analysis

Kazuki Kurimoto: Investigation, Methodology, Formal analysis, Writing – review and editing

Yasuhiko Saito: Investigation, Methodology, Formal analysis, Writing – review and editing Yasuto Kunii: Investigation, Methodology, Formal analysis, Writing – review and editing Ken-ichiro Kubo: Investigation, Methodology, Formal analysis, Writing – review and editing Hiroaki Wake: Investigation, Methodology, Formal analysis, Writing – review and editing Taku Nagai: Investigation, Methodology, Formal analysis, Writing – review and editing Manabu Makinodan: Conceptualization, Methodology, Supervision, Project administration, Funding acquisition, Writing – original draft, Writing – review and editing.

## Data Availability

The data supporting the findings of this study are available from the corresponding author upon reasonable request.

---

**Figure 1—figure supplement 1.**
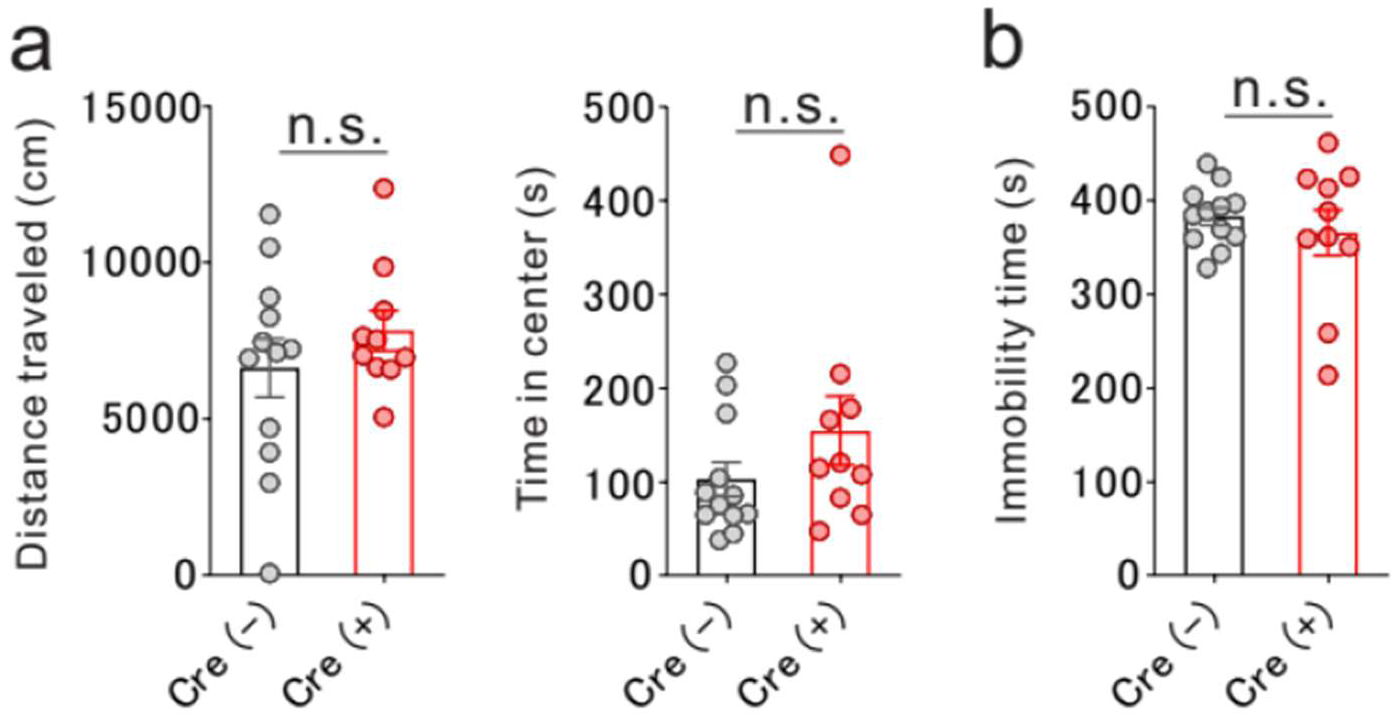
Open-field and tail-suspension tests in mice with *Myrf* deletion in the mPFC. (a) Open-field test showing total distance traveled (left) and time spent in the center area (right) in Cre⁻ and Cre⁺ mice. No significant differences were observed between the groups (distance traveled: two-tailed unpaired t-test, p=0.5249; time in center: two-tailed unpaired t-test, p=0.1970), showing no detectable group differences in general locomotor activity or anxiety-like behavior. (b) Tail-suspension test showing immobility time in Cre⁻ and Cre⁺ mice. No significant difference was detected (two-tailed unpaired t-test, p=0.4879), showing no detectable group difference in depression-like behavior. Data are presented as mean ± standard error of the mean (SEM). Each dot represents an individual animal (Cre⁻, n=12; Cre⁺, n=10). n.s., not significant.

**Figure 2—figure supplement 1.**
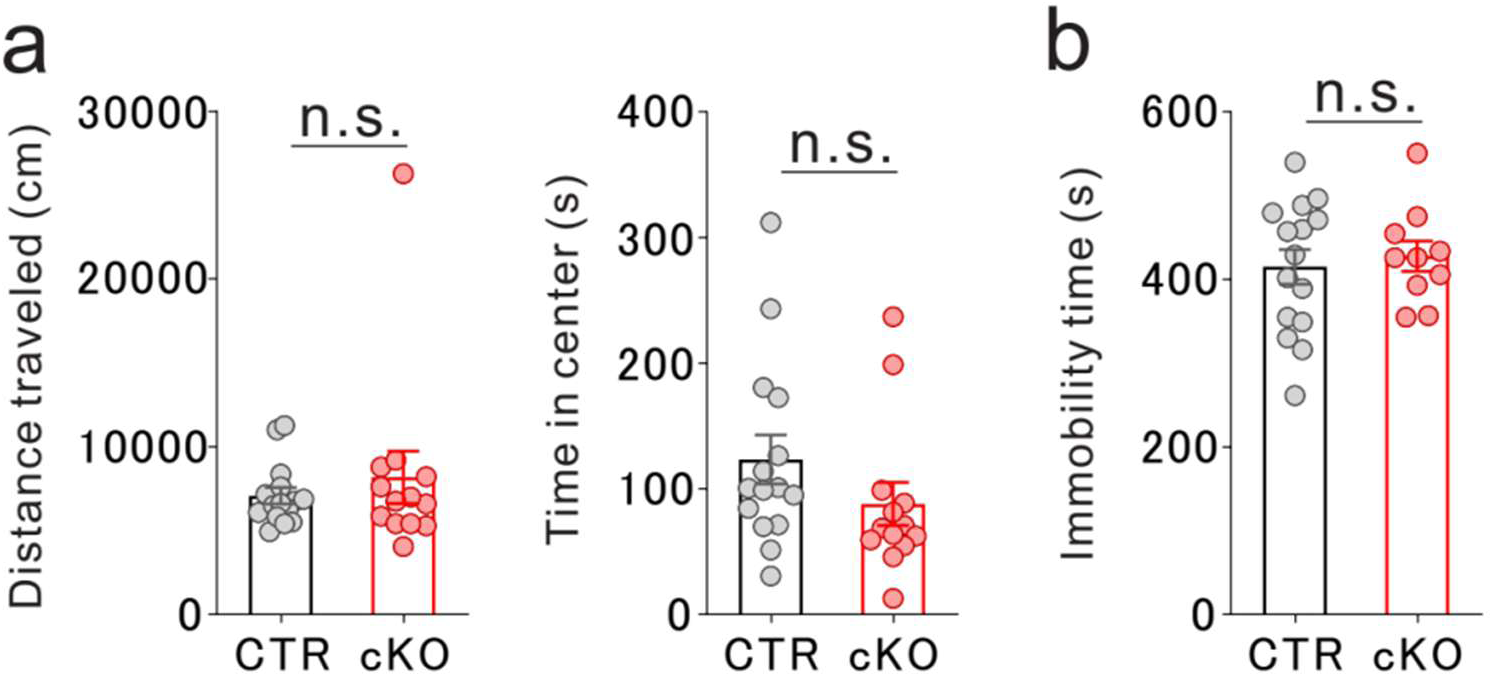
Open-field and tail-suspension tests in mice with developmental *Myrf* deletion in Olig2-lineage cells. (a) Open-field test showing total distance traveled (left) and time spent in the center area (right) in CTR and cKO mice. No significant differences were observed between the groups (distance traveled: two-tailed unpaired t-test, p=0.4804; time in center: two-tailed unpaired t-test, p=0.1879; CTR, n=15; cKO, n=13), showing no detectable group differences in general locomotor activity or anxiety-like behavior. (b) Tail-suspension test showing immobility time in CTR and cKO mice. No significant difference was detected (two-tailed unpaired t-test, p=0.6675; CTR, n=15; cKO, n=10), showing no detectable group difference in depression-like behavior. Data are presented as mean ± standard error of the mean (SEM). n.s., not significant.

**Figure 3—figure supplement 1.**
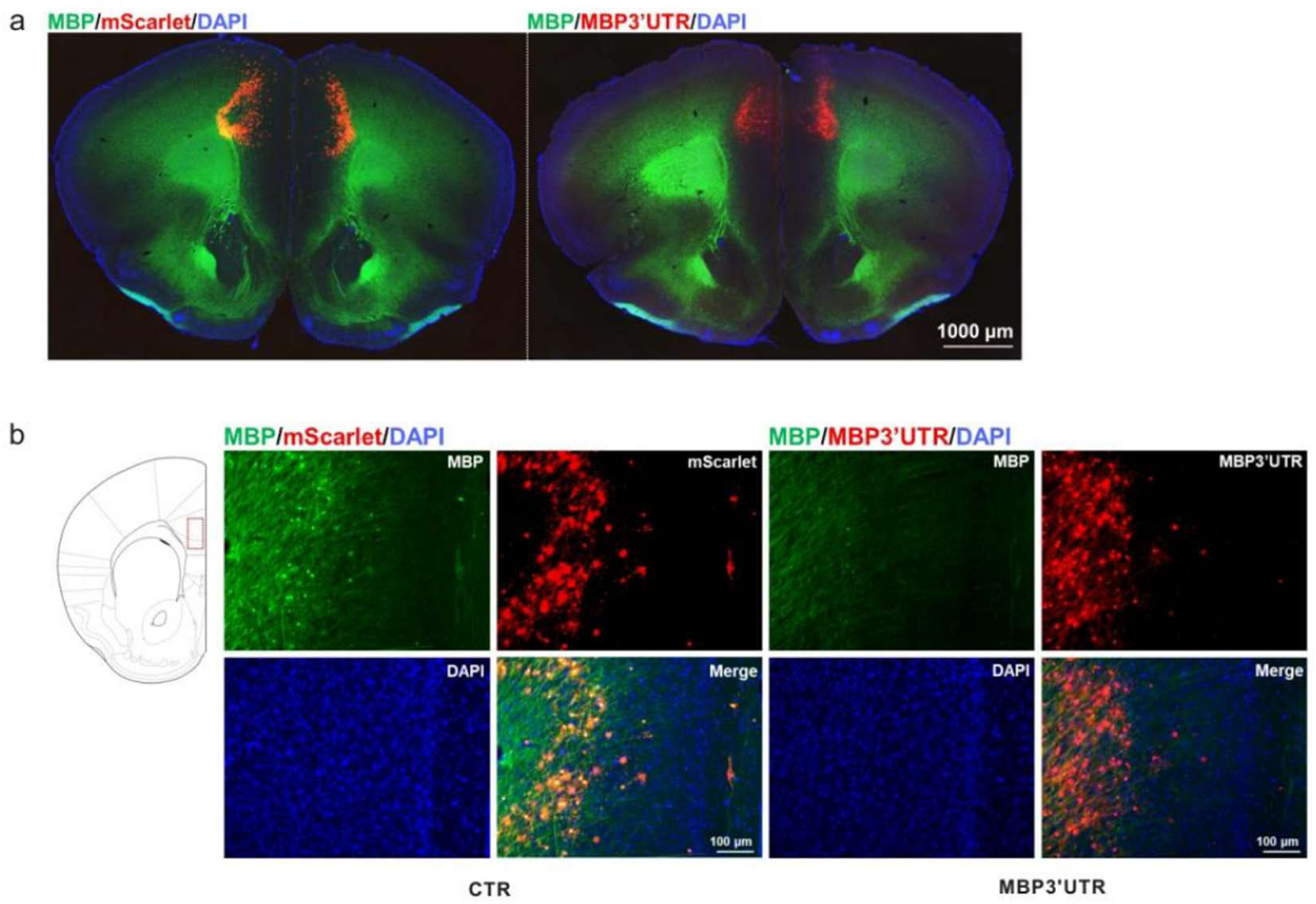
Reduction of cortical myelination following MBP3’UTR expression. (a) Representative coronal sections showing myelin basic protein (MBP) immunostaining in control adeno-associated virus (AAV; mScarlet) and MBP3’UTR-injected mice. MBP (green), mScarlet or MBP3’UTR expression (red), and 4′,6-diamidino-2-phenylindole (DAPI) (blue) are shown. MBP3’UTR-injected mice exhibit a marked reduction in MBP immunoreactivity across the cortex. Scale bar, 1000 µm. (b) Higher-magnification images of the medial prefrontal cortex (mPFC) (layers I–V). MBP immunoreactivity is substantially decreased in MBP3’UTR-injected mice compared with controls. Scale bar, 100 µm.

**Figure 3—figure supplement 2.**
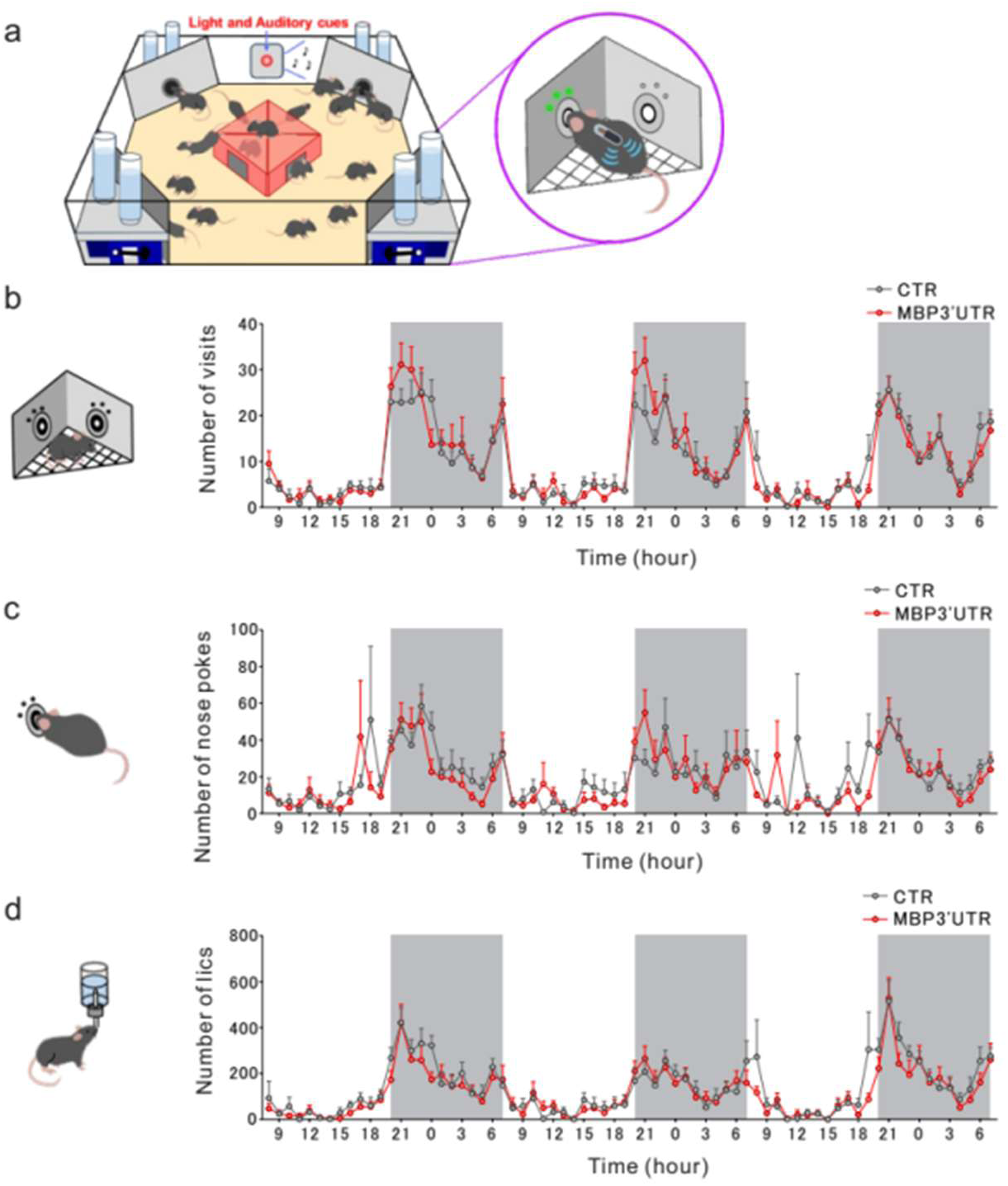
IntelliCage assessment of baseline activity under group-housed conditions. (a) Schematic of the IntelliCage system used for automated behavioral monitoring in group-housed mice. The setup allows continuous tracking of individual animals, including corner visits, nose pokes, and licking behavior, enabling assessment of spontaneous activity and social behaviors. (b–d) Circadian activity profiles across 24-h cycles during the adaptation and training phases. Hourly counts of (b) corner visits, (c) nose pokes, and (d) licks are shown for control and MBP3’UTR-injected mice. Both groups exhibited comparable diurnal patterns, with increased activity during the dark phase and reduced activity during the light phase. No significant differences were observed between the groups, showing no detectable group differences in baseline locomotor activity, motivation, or task acquisition.

**Figure 5—figure supplement 1.**
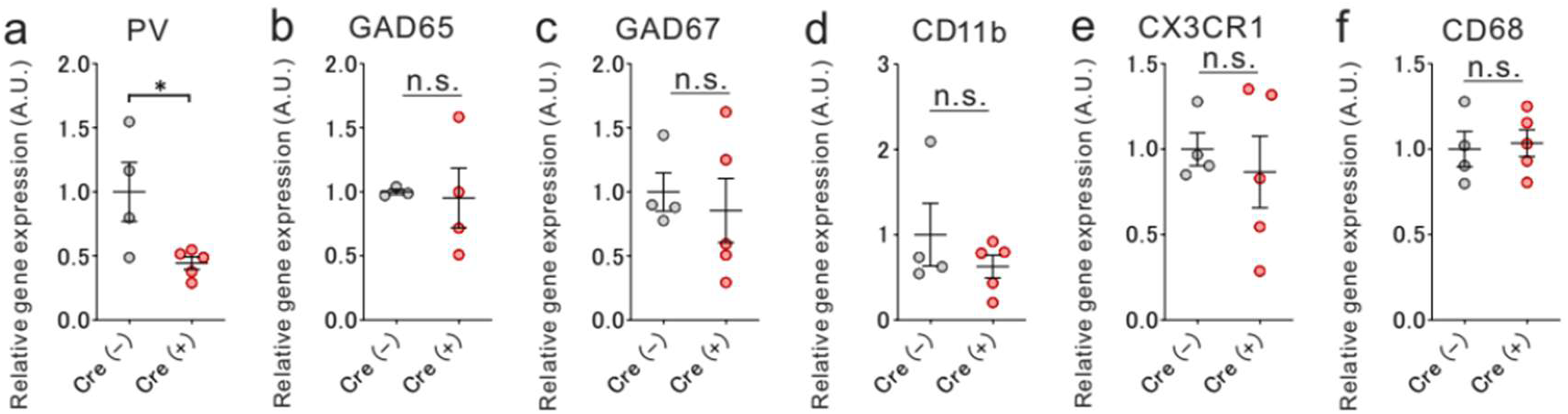
Selective reduction of PV expression without evidence of overt microglial activation in hypomyelinated mPFC. (a–c) qRT–PCR analysis of interneuron-related gene expression in the mPFC of AAV-mediated *Myrf* deletion (Cre⁺) and control (Cre⁻) mice. (a) Relative expression of PV mRNA is reduced in hypomyelinated mice (two-tailed unpaired t-test, p=0.0321; Cre⁻, n=4; Cre⁺, n=5). In contrast, expression levels of GABA-synthesizing enzymes (b) GAD65 (two-tailed unpaired t-test, p=0.8702; Cre⁻, n=3; Cre⁺, n=4) and (c) GAD67 (two-tailed unpaired t-test, p=0.6553; Cre⁻, n=4; Cre⁺, n=5) show no significant differences between the groups, suggesting a preferential effect on PV expression. (d–f) qRT–PCR analysis of microglial marker genes in the mPFC. Relative expression of (d) CD11b (two-tailed unpaired t-test, p=0.3295; Cre⁻, n=4; Cre⁺, n=5), (e) CX3CR1 (two-tailed unpaired t-test, p=0.6128; Cre⁻, n=4; Cre⁺, n=5) and (f) CD68 (two-tailed unpaired t-test, p=0.79471; Cre⁻, n=4; Cre⁺, n=5) show no significant differences between Cre⁺ and Cre⁻ mice, providing no evidence of overt microglial activation based on these markers following developmental hypomyelination. Data are presented as mean ± standard error of the mean (SEM). n.s., not significant.

**Supplementary Table 1.**
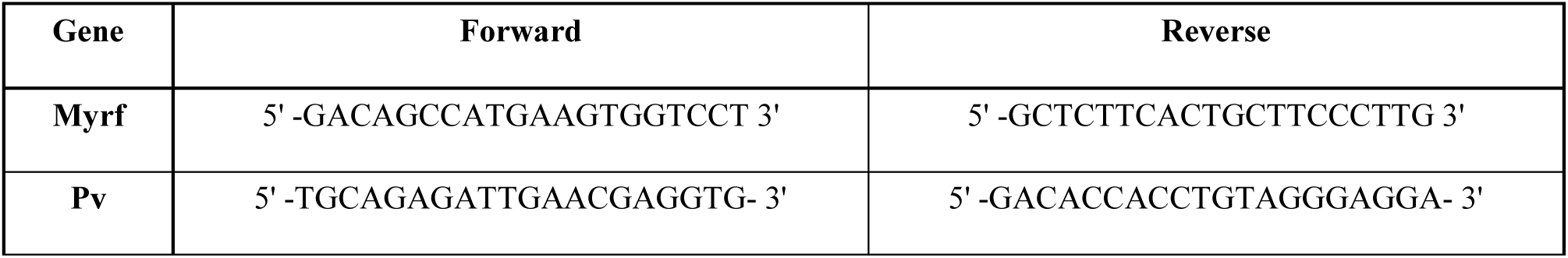
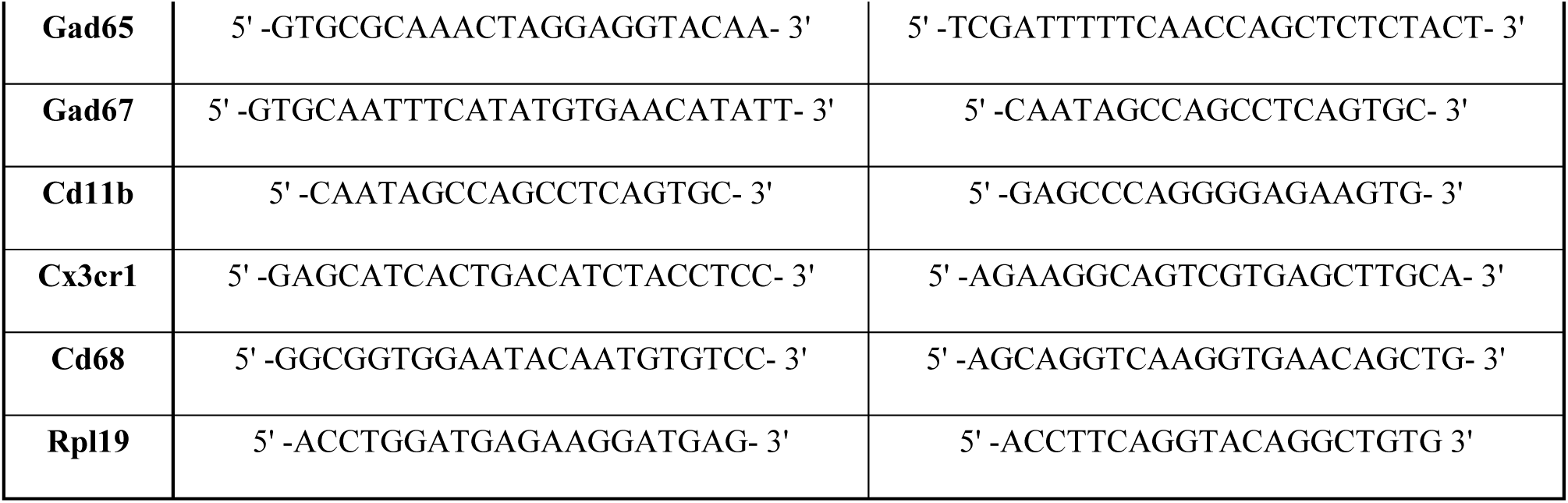
Primer sequences used for quantitative RT–PCR analysis.

